# Regulators of TNFα Mediated Insulin Resistance Elucidated by Quantitative Proteomics

**DOI:** 10.1101/2020.06.22.165472

**Authors:** Rodrigo Mohallem, Uma K. Aryal

## Abstract

Obesity is a growing epidemic worldwide and is a major risk factor for several chronic diseases, including diabetes, kidney disease, heart disease, and cancer. Obesity often leads to type 2 diabetes mellitus (T2DM, via the increased production of proinflammatory cytokines such as tumor necrosis factor-α (TNFα). Our study combines different proteomic techniques to investigate the changes in the global proteome, secretome and phosphoproteome of adipocytes under chronic inflammation condition, as well as fundamental cross-talks between different cellular pathways regulated by chronic TNFα exposure. Our results show that many key regulator proteins of the canonical and non-canonical NF-κB pathways, such as Nfkb2, and its downstream effectors, including Csf-1 and Lgals3bp, directly involved in leukocyte migration and invasion, were significantly upregulated at the intra and extracellular levels, culminating in the progression of inflammation. Our data provides evidence of several key proteins that play a role in the development of insulin resistance.

## Introduction

The steep rise in obesity in the last few decades has become a topic of growing concern worldwide. In the United States alone, obesity has increased by an alarming 9% during the last twenty years. It is currently estimated that 39.8% of the adult American population is obese, with an additional third that is overweight (Hales, 2017; Hales et al., 2018). Severe obesity is associated with an increased risk of development of various ailments, among which insulin resistance is particularly concerning (Reaven, 1988).

In obese patients, insulin resistance is often accompanied by dysfunctional pancreatic islet β-cells, characterized by a decreased secretory capacity. Insulin secretion by dysfunctional islet β-cells becomes insufficient to overcome the loss in cellular insulin sensitivity, culminating in hyperglycemia and type-2 diabetes mellitus (T2DM) (Kahn, 2001). Furthermore, obesity induced adipose tissue hyperplasia and hypertrophy is often associated with an increased secretion of adipose tissue-derived cytokines, commonly referred as adipokines (Jo et al., 2009). Increased production of pro-inflammatory adipokines, such as tumor necrosis factor-α (TNFα), interleukine-6 (IL-6), and interleukine-3 (IL-3), induces insulin resistance, either by directly disrupting the canonical insulin signaling pathway, or by stimulating the activation of additional inflammatory pathways (Tilg and Moschen, 2008). In particular, TNFα has been shown to be overexpressed in obese mice in both transcriptional and translational levels. Abolishment of TNFα sensitivity, in turn, was shown to promote increased insulin response and consequent increased glucose uptake in obese rats (Hotamisligil et al., 1993). Similar reports indicate that TNFα is also overexpressed in human adipose tissue and muscle tissue of obese patients, relative to lean patients (Kern et al., 1995; Saghizadeh et al., 1996).

The abnormal secretion of adipokines induces the recruitment and invasion of B cells, T cells, and subsequently macrophages to the adipose tissue. After infiltration, IFN-γ and IL-17 secreting T-cells stimulate the proinflammatory activation of adipocyte tissue macrophages (ATM’s) (Sell et al., 2012). Necrotic white adipose tissue, a hallmark of obesity, is also a major driver for macrophage infiltration. Over 90% of ATMs have been shown to accumulate and surround dead adipocytes (Cinti et al., 2005), forming a crown-like structure that is associated with increased TNFα and IL-6 secretion (Strissel et al., 2007). Additionally, adipocytes release free fatty acids, which interacts with ATM Toll-Like Receptor 4 (TLR4), triggering the activation of the pro-inflammatory Nuclear Factor-κB (NF-κB) pathway. Subsequent to the activation of the NF-κB cascade, the secretion of other adipokines are further stimulated, notably monocyte chemotactic protein-1 (MCP-1), a cytokine that promotes the recruitment of leukocytes to the inflammation site (Bai and Sun, 2015).

While the regulation of insulin responses and development of the pathophysiology of insulin resistance due to overexpression of TNFα and other pro-inflammatory cytokines is well documented, the functional links and interactions that mediate insulin resistance are complex and largely unexplored. Identifying the primary effectors and elucidating crosstalks between underlying molecular mechanisms involved in disrupting insulin signaling and other inflammatory pathways by TNFα overexpression, is a crucial step for a comprehensive understanding of T2DM development. In this study, we focused on a holistic approach to identify pivotal adipocyte proteins that are significantly altered upon chronic TNFα exposure. Through a combination of global proteomics, secretomics and phosphoproteomics our data revealed several effector proteins involved in the progression of inflammation, significantly regulated during prolonged TNFα exposure. Our data unveils novel candidate proteins that could directly modulate the physiological signaling cascade of canonical and noncanonical pathways in response to TNFα. These results present valuable insight on the dynamic changes in the proteome landscape of TNFα chronically adipocytes, and provides an invaluable resource for future investigations on potential therapeutic targets for insulin resistance and T2DM.

## Results

### Integrated Label-Free Quantitative Proteomic Analysis

We utilized an integrated label-free quantitative global proteomic, secretomic and phosphoproteomic approach to shed light on the complex regulations at the proteome level of adipocytes chronically exposed to TNFα. Murine 3T3-L1 pre-adipocytes, which have become fundamental in metabolic disease researches, were chemically induced to differentiate, using a standard protocol (ATCC). After full differentiation, inflammation was replicated in-vitro by chronically treating adipocytes with 10 nM insulin and 2 ng/mL TNFα. As a positive control, adipocytes were chronically treated with 10 nM insulin alone. Previous reports have suggested that chronic TNFα treatment for a period of 4 days is sufficient to induce insulin resistance in both myocytes and murine adipocytes (Hoehn et al., 2009; Yoon et al., 2011). To gain an insight on the dynamic effects of TNFα-induced insulin resistance, we performed a time-resolved proteomic analysis, at 4 days (4D) and 8 days (8D) of chronic treatment (Figure 1A). Spent media was collected and used for secretome analysis (Figure S1A). Adipocytes were then collected and efficiently lysed with a homogenizer. Total protein was extracted, cysteine disulfide bonds were reduced and alkylated, followed by proteolysis with Lys-C/Trypsin, as previously described (Hedrick at al., 2015). Peptides were desalted and 1 ug was used for global proteomic analysis (Figure 1B). LC-MS/MS data were searched against Uniprot Mus musculus database using MaxQuant platform (Cox and Mann, 2008; Tyanova et al., 2016), and subsequent statistical analysis was performed with the Perseus bioinformatics software (Tyanova et al., 2016) (Figure 1C).

**Figure 1.**
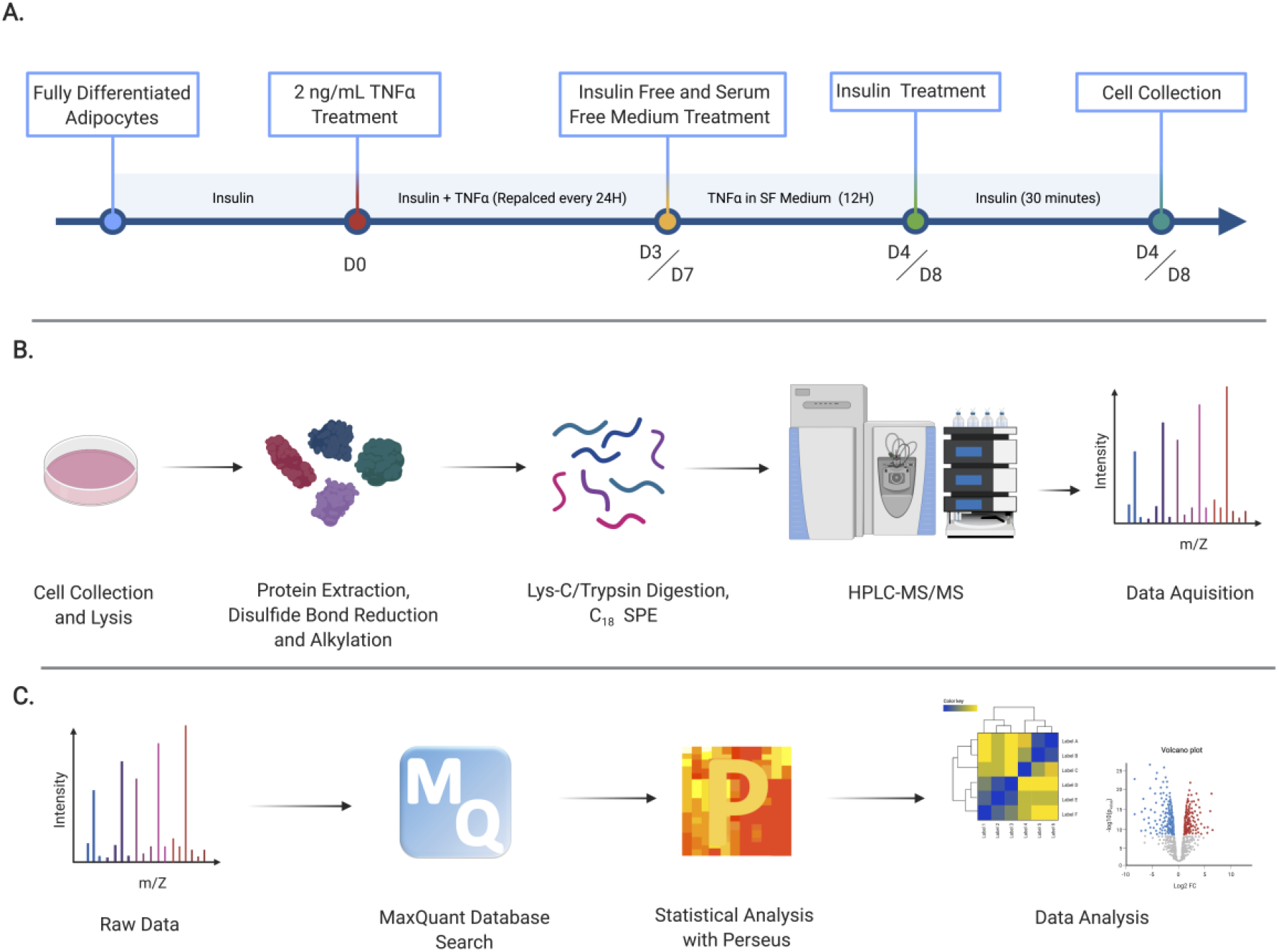
Experimental design and label-free quantitative global proteomic analysis workflow. (A) 3T3-L1 fully differentiated murine adipocytes were chronically treated with 10 nM insulin, and experimental group were treated with an additional 2ng/mL TNFα for 4 or 8 consecutive days. 12 hours prior to cell collection, culture medium was replaced with serum free and insulin free medium, fol lowed by a 30-minute insulin treatment before consecutive cell collection. (B) Total protein was extracted from the cell lysate, with sequential reduction and alkylation of cysteine disulfide bonds. Proteins were digested with Lys-C/Trypsin, desalted and analyzed by HPLC-MS/MS. (C) Raw LC-MS/MS data were searched using the MaxQuant platform. Perseus was used for statistical analysis. Filtered, log-transformed and normalized data were used for subsequent analysis and visualization.

We identified more than 24500 peptides, mapped to over 2992 proteins. From the total proteins identified, 2368 proteins were quantified (LFQ > 0) in at least 2 biological replicates from the same treatment group. Our dataset ranks first in combined global proteomics, phosphoproteomics and secretomics to characterize the dynamics of protein expression in murine adipocytes during chronic TNFα exposure.

### Chronic TNFα-Treatment Induces Extensive Regulations in the Proteome of 3T3-L1 Adipocytes

The heatmap representation of the Z-transformed LFQ values from 2369 proteins identified shows a very clear and consistent distribution of protein expression patterns between treatment groups, on both 4D and 8D timepoints (Figure 2A). The consistency between each biological replicate was further evidenced by Pearson correlation analysis. As indicated by the color scale on the correlation plots, replicates from the same treatment group show distinct correlation values than those across treatments, indicating the high consistency and reproducibility of the MS/MS approach (Figure 2B). The consistency and similarity between biological replicates are further confirmed by principal-component analysis, in which a distinct separation and aggregation between different treatments is clearly observed (Figure 2SA, B).

**Figure 2.**
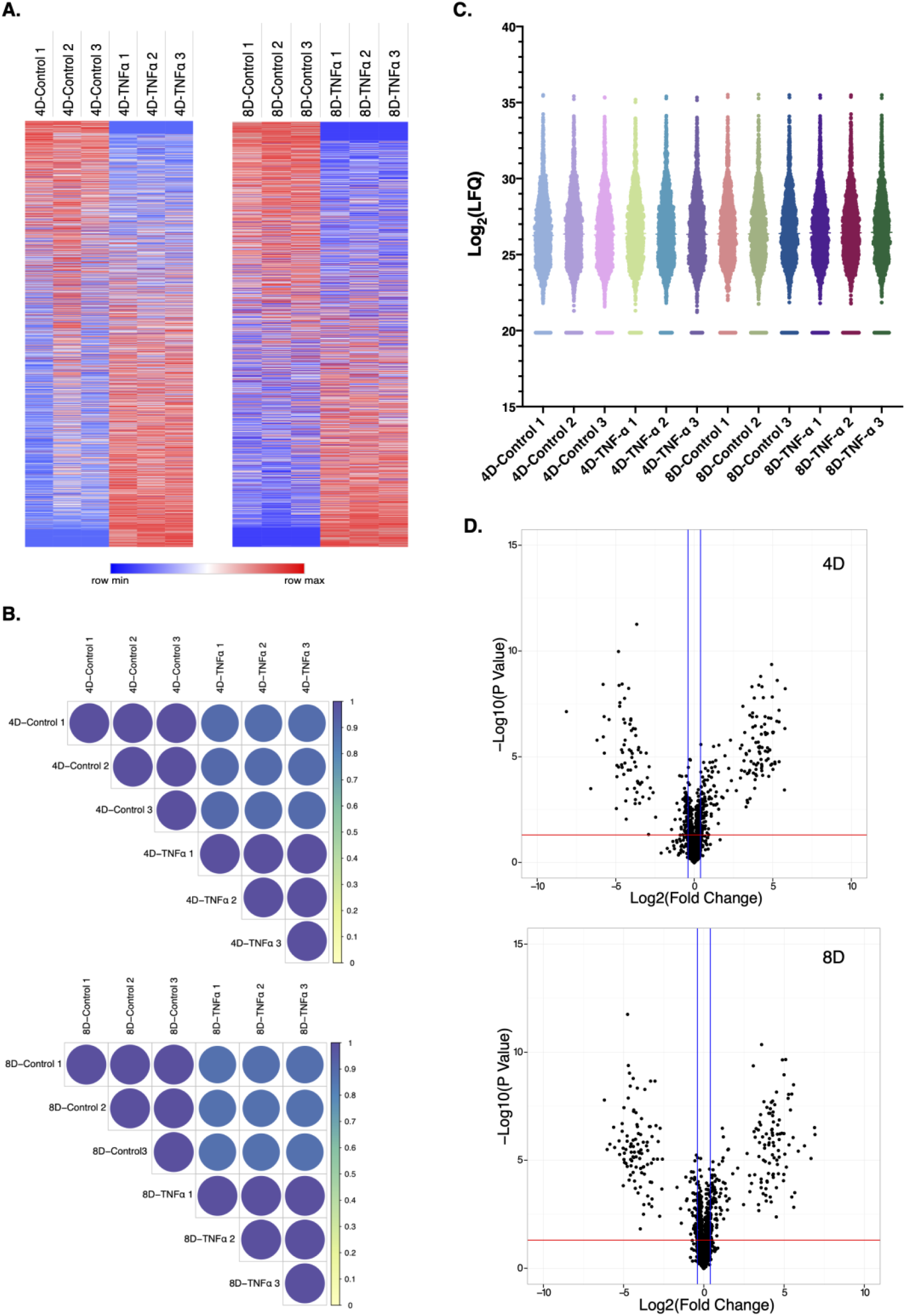
Global proteomic analysis of TNFα chronically treated adipocytes. (A) Heatmap of all 2369 quantified proteins for both 4D and 8D timepoints. Color scale represents Z-scored LFQ values. (B) Pearson correlation coefficient of biological replicates from each treatment group. Color scale indicates Pearson correlation coefficient. (C) Violin plot of Log-transformed LFQ values from all analyzed samples. Colors were arbitrarily assigned to each biological replicate. (D) Volcano plots of all quantified proteins in each timepoint. Horizontal red line represents the Log_10_ (p value) cutoff, and vertical blue lines represent the Log_2_ (Fold Change) cutoff.

Moreover, violin plot suggests the remarkably consistent distribution of log transformed LFQ values and probability density across all samples, with values clustering within the first and third interquartile ranges. All biological replicates showed a normal distribution-like shape, with uniform mean values (Figure 2C).

To study the changes in protein expression induced by chronic inflammation we aimed to identify all proteins, which were significantly upregulated or downregulated in TNFα treated cells compared to the control for each time point. We selected all proteins that were significantly different between the TNFα treated and control groups (p < 0.05, Two-tailed Students T-test), with a Log_2_(Fold Change) ≥ |0.38|. From the 2369 quantified proteins, a total of 347 and 340 proteins met the criteria for 4D and 8D timepoints, respectively. These proteins can be distinguished in the volcano plot analysis, in which proteins represented above the cutoff lines are significantly regulated (Figure 2D). It is interesting to notice the very symmetric distribution of the Volcano plot, indicating that chronic TNFα treatment induces a complex regulation of the proteome of 3T3-L1 adipocytes, inducing both upregulation and downregulation of several different proteins.

### Differentially Expressed Proteins Show Highly Consistent Expression Levels

For the past decade, the identification of molecular links between obesity, inflammation and insulin resistance has become a major focus of discussion. It has become clear that the development of insulin resistance during obesity is resultant of a complex and interconnected underlying pathobiochemistry (Yazıcı and Sezer, 2017). However, few reports have focused on an omics perspective to dissect the large-scale progression of insulin resistance. Although proteomic analysis has been previously performed on adipocytes (Chan et al., 2019), there are no reports to date that investigate the effects of prolonged inflammation on a proteome scale. To unveil the dynamics of the cellular proteome during chronic TNFα exposure, we selected all quantified proteins mapped for each treatment condition, filtering for identified proteins with an MS/MS count ≥ 2 in at least 2 biological replicates to increase data accuracy, and performed a direct comparison. To visualize unique and common proteins found in each treatment and time-point, we performed a circular plot analysis, in which a string connects overlapping proteins. The majority of mapped proteins were common to at least two different treatments, with expected greater overlap between treatment groups from the same time-point (Figure 3A). We then set to identify proteins that were only significantly regulated during a specific timepoint, and proteins that were continuously regulated between the 4D and 8D groups. We found 166 proteins uniquely identified at 4D, 130 uniquely identified at 8D, and 119 proteins that were significantly regulated at both 4D and 8D (Figure 3B). It is worth noting that the majority of the proteins which only show significant regulation at one specific timepoint appear on the total quantified protein list of both timepoints, likely due to a more temporal regulation at different TNFα exposure times.

**Figure 3.**
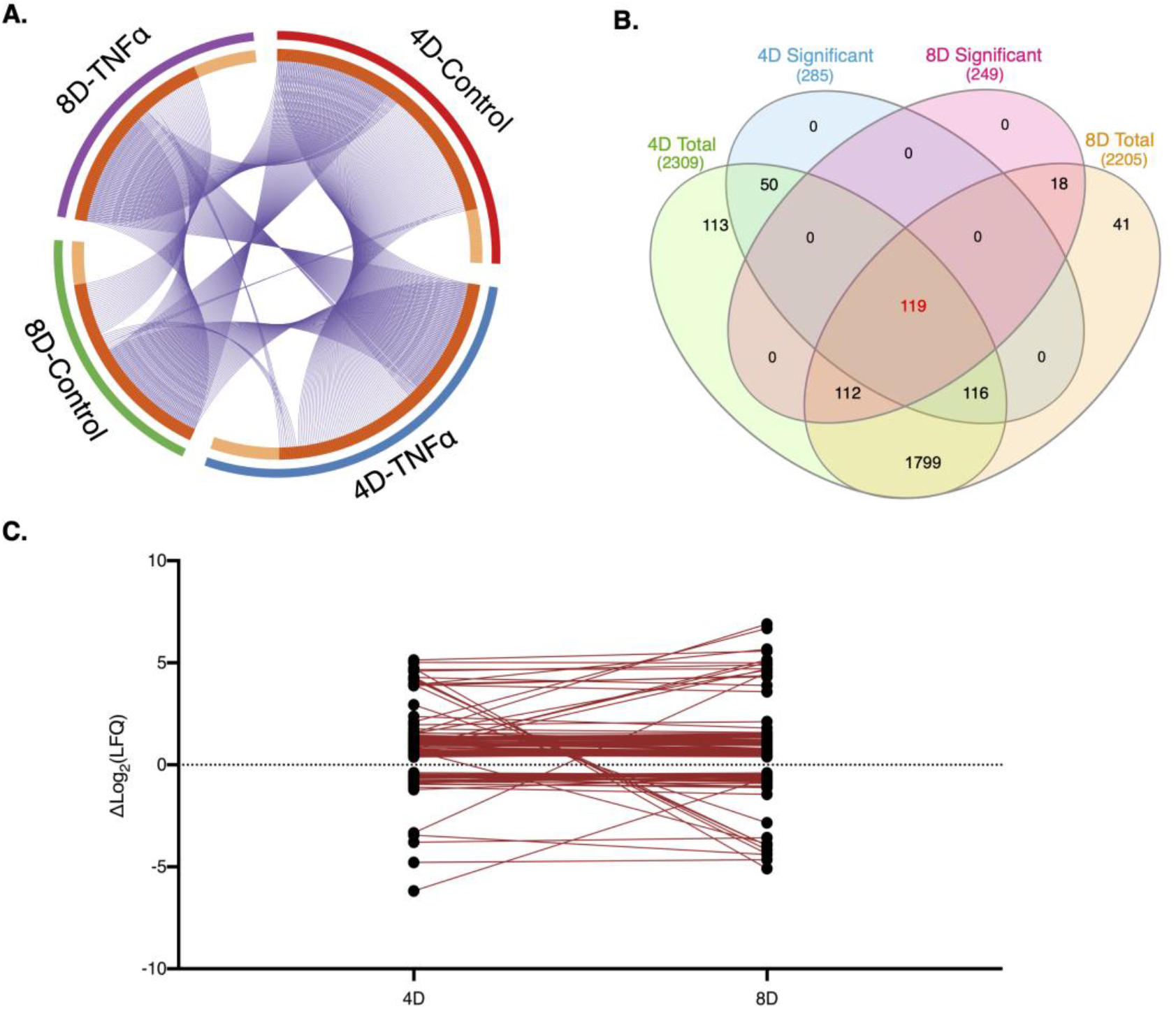
Comparative analysis 4D and 8D. (A) Circular plot representation of proteins identified in each treatment group. Light orange indicates uniquely mapped proteins. Blue strings indicate proteins common to two or more treatment groups. (B) Venn Diagram i llustrating the overlap between significantly regulated proteins at 4D (blue) and 8D (yellow) timepoints and total quantified proteins at 4D (green) and 8D (orange). (C) Line plot depicting the variance in the log-transformed fold-change values of the overlapping 119 proteins between 4D and 8D timepoints.

The pathogenesis of insulin resistance is directly associated with chronic low-grade inflammation during obesity, which, in turn, promotes gradual and sustained changes on the molecular landscape of the adipose tissue (Chen et al., 2015). By characterizing the proteins that were consistently significantly regulated during prolonged TNFα treatment, we were able to gain a fundamental insight on the key proteins that are continuously regulated during the chronic treatment, while also allowing for a crucial analysis of the changes in expression levels as the exposure time to TNFα increases. Therefore, we hereon focused our analysis on the significantly regulated proteins continuously expressed on both 4D and 8D timepoints.

Aiming to investigate how the expression levels of the identified proteins changed between timepoints, we plotted and linked the expression levels of each individual protein at 4D and at 8D. From the 119 proteins identified, 94 showed a very consistent trend in log-transformed fold change values across the two timepoints, with a difference between values at 4D and 8D of 0.38 or less. Conversely, 17 proteins showed a greater difference in fold change values. It is interesting to note that 8 proteins were observed to invert regulation patterns, that is, switching from upregulated at 4D to downregulated at 8D, or vice versa (Figure 3C).

To categorize and visualize the regulation patterns of each protein, we categorized the continuously expressed proteins into three distinct groups, upregulated, downregulated and reversed. Heatmaps were then generated for each group, highlighting the differences in Z-score for each protein across all biological replicates and timepoints. The expression patterns for both downregulated and upregulated groups was highly consistent across all replicates, on both 4D and 8D timepoints (Figures 4A and 4B). The number of upregulated proteins were noticeably higher, and most were directly involved in inflammatory processes. We identified several proteins that are activated by, and act as downstream effectors in pro-inflammatory signaling pathways. Notably, nuclear factor NF-κB p100/p52 subunit (Nfkb2), which is a precursor for transcriptional activator complex p52/RelB/NF-κB in the non-canonical TNFα induced signaling pathway (Liu et al., 2017), and its activation has implications in the progression of metabolic inflammation (Choudhary et al., 2011). We also identified several different proteins involved in the IFNα/β signaling pathway, including the transcription factors Signal transducer and activator of transcription 1 (Stat1) and 2 (Stat2), Interferon-inducible GTPase 1 (Iigp1), and interferon-induced protein with tetratricopeptide repeats 2 (Ifit2) and 3 (Ifit3), suggesting a potential cross-talk between the TNFα and IFNα/β activated pathways. Our data also confirms the overexpression of cytokines directly correlated with macrophage recruitment, particularly macrophage migration inhibitory factor (Mif) (Hirokawa et al., 1998). Conversely, downregulated proteins included metabolic proteins such as alcohol dehydrogenase 1 (Adh1), arylsulfatase A (Arsa) and B (Arsb), and acid ceramidase (Asah1).

**Figure 4.**
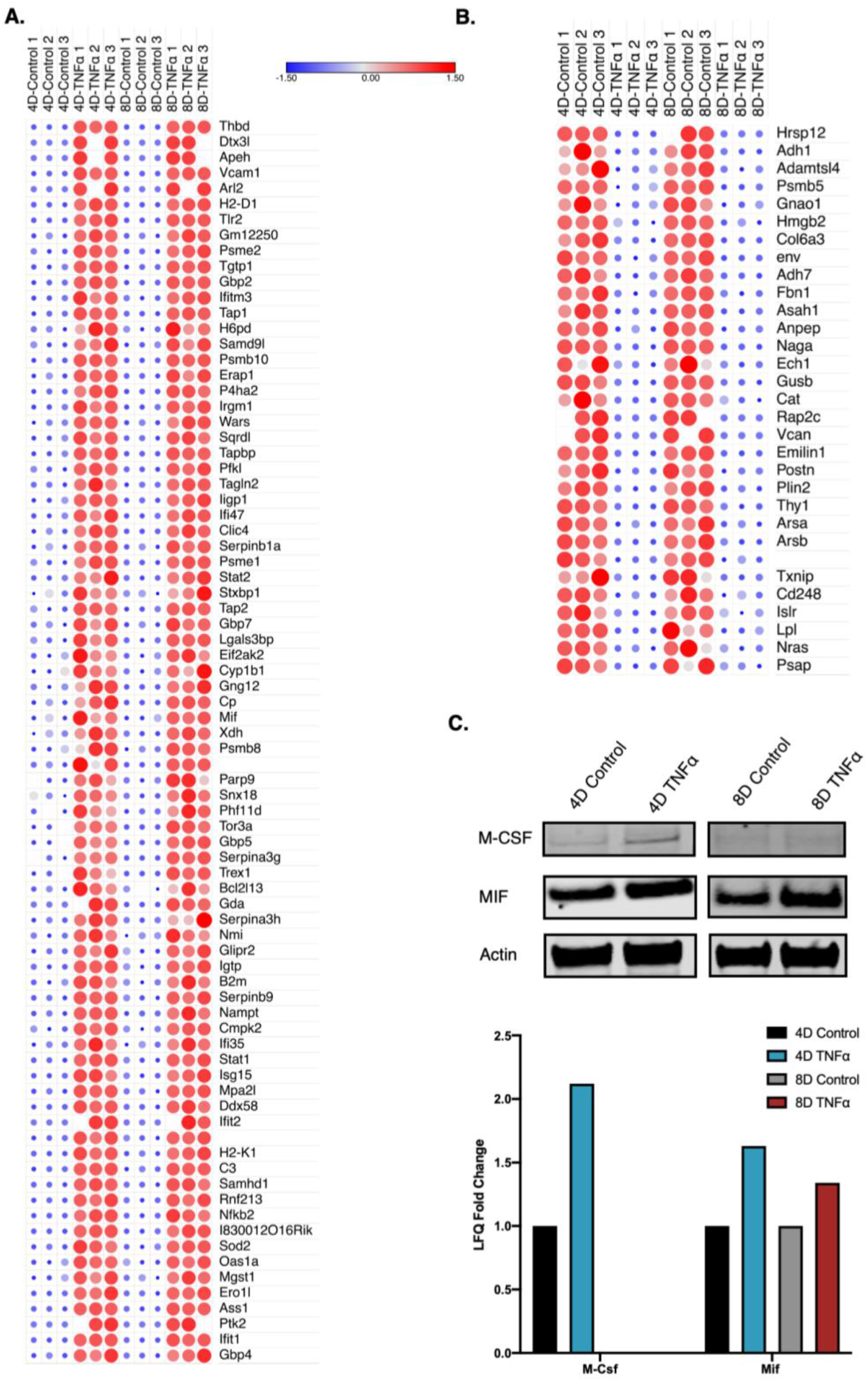
Differentially expressed proteins are mostly upregulated. (A) Heat-map depiction of upregulated continuously expressed proteins between 4D and 8D timepoints. Color scale and relative dot size represents Z-scored LFQ values. Blank spaces represent missing values. (B) Heat-map depiction of downregulated continuously proteins between 4D and 8D timepoints. Color scale and relative dot size represents Z-scored LFQ values. Blank spaces represent missing values.(C) Western blot analysis of M-Csf and Mif expression levels in 4D and 8D TNFα treated adipocytes and corresponding LFQ fold change.

We then selected one of the proteins discussed above, Mif1, and performed western blot analysis, to validate the quantitative information acquired through the proteomic analysis. Consistent with the proteomics results, Mif1 shows a consistent increased signal at both 4D and 8D across all biological replicates from the TNFα treated group, compared with the control. In addition to Asah1 and Mif1, we probed for the cytokine Colony-stimulating factor 1 (Csf1), which has been shown to regulate the differentiation, proliferation, survival and activation of monocytes and macrophages (Chang et al., 2014). Our proteomics results indicate that Csf-1 had a 1.37 fold increase in its expression levels in TNFα treated cells at 4D, but it was not identified at 8D. Western blot analysis clearly showed Csf1 was upregulated at 4D of TNFα treatment, and no signal was observed at 8D, which is very consistent with the MS/MS data (Figure 4C). These data diverge from previous studies, which reported no significant changes in Csf1 gene expression in the adipose tissue of obese mice (Sugita et al., 2007). Our results, conversely, suggest that TNFα induced chronic inflammation promotes an early overexpression of Csf1 in response to TNFα, which likely contributes to the invasion of leukocytes to the adipose tissue.

### Continuously Regulated Proteins in 4D and 8D Mediate insulin Resistance

To gain a better perspective of the functional biological process that accompanied chronic TNFα treatment, we performed Gene Ontology (GO) enrichment analysis (Ashburner et al., 2000; The Gene Ontology Consortium, 2019), and annotated the significant (p < 0.05) terms to a network analysis of all continuously expressed up and downregulated proteins that had consistent expression levels across 4D and 8D of treatment, with a variation in Log_2_(LFQ) ≤ |0.38|. We have found that “ceramide catabolic process” was the most significantly downregulated biological process, followed by response to nutrient (Figures 5A). The release and accumulation of free fatty acids in the adipose and muscle tissue, specifically the sphingolipid ceramide, has been directly associated with the progression of insulin resistance in obese mice (Sokolowska and Blachnio-Zabielska, 2019). Our results suggest that chronic exposure to TNFα induces a decrease in the levels of metabolic enzymes that partake in ceramide metabolism, namely Asah1, prosaposin (Psap), and alpha-N-acetylgalactosaminidase (Naga), and likely promotes an increase in the intra and extracellular concentrations of ceramides.

**Figure 5.**
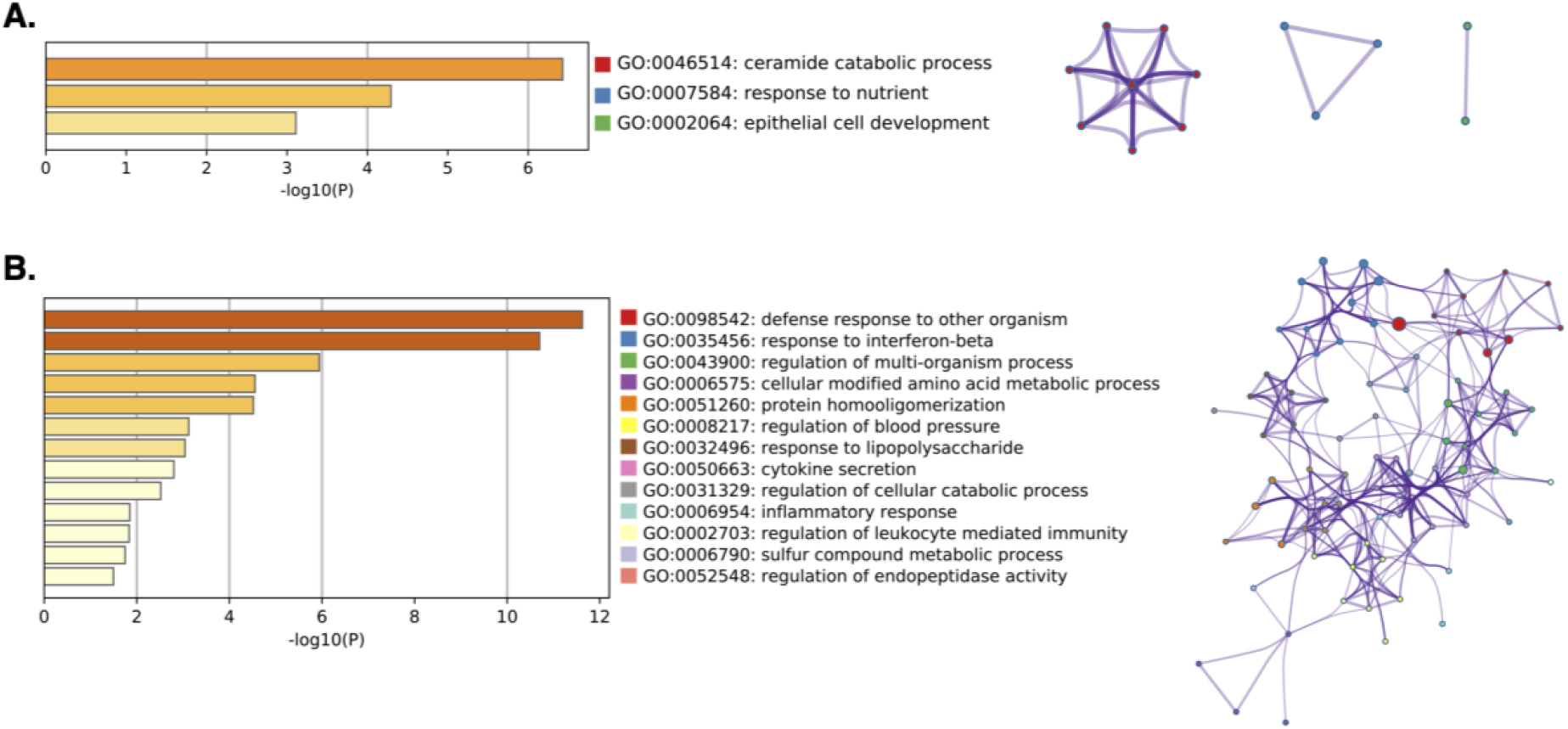
Biological processes regulated by chronic TNFα treatment. (A) Significantly downregulated GO Biological Processes for continuously expressed proteins in TNFα treated cells. Network plot depicts the relationship between enriched terms. Nodes represent individual enriched terms and are colored based on its cluster ID. Terms with similarity >0.3 are connected by strings. (B) Significantly upregulated GO Biological Processes for continuously expressed proteins in TNFα treated cells. Network plot depicts the relationship between enriched terms. Nodes represent individual enriched terms and are colored based on its cluster ID. Terms with similarity >0.3 are connected by strings.

Our results also show significant upregulation in pro-inflammatory and cell mediated immune response pathways. The upregulated biological process “response to interferon beta”, “response to lipopolysaccharide”, “cytokine secretion”, “inflammatory response” and “regulation of leukocyte mediated immunity” are particularly interesting (Figures 5B). All upregulated gene ontology terms were also shown to be tightly linked, which suggests a close overlap among the proteins mapped to each biological process.

### Chronic TNFα Treatment Promotes Differential Regulation of Secreted Proteins

The adipose tissue is often associated with the storage of triacylglycerol and other fatty acids. Although primarily a site for nutrient storage, the adipocyte tissue also plays important endocrine, paracrine and autocrine roles, mediated by the secretion of several different hormones and adipokines, and numerous other growth factors, enzymes, complement factors and matrix proteins (Coelho et al., 2013). Obesity promotes a drastic change in the secretion profile of the adipose tissue, marked by the release of proinflammatory cytokines, including TNFα and IL-6, which can directly regulate inflammation, and, by extent, insulin resistance (Makki et al., 2013). Despite the extensive characterization of proinflammatory cytokines and their roles in insulin resistance, the secretome of adipocytes during inflammation remains unexplored. To gain a wholistic perspective on the dynamics of intracellular and secreted protein levels during chronic inflammation, we combined our global proteome with a parallel secretome analysis of adipocytes chronically treated with TNFα. Briefly, prior to cell scraping, spent media was collected and subsequently concentrated with a Vivaspin 6™ Centrifugal Concentrator 10,000 MWCO PES. 50 ug of concentrated sample were reduced and alkylated, followed by proteolysis with Lys-C/Trypsin, before LC-MS/MS analysis. Raw data was analyzed as described previously.

We identified a total of 2149 peptides, mapped to 269 protein groups at 4D, and 1929 peptides, assigned to 243 protein groups at 8D of TNFα treatment. From the total proteins identified, 125 and 118 proteins were quantified in at least 2 biological replicates, at 4D and 8D, respectively. The expression patterns between the total quantified secreted proteins were highly consistent among replicates, and across timepoints, as evidenced by the heatmap representation of Z-scored LFQ values (Figure S2A). To study the dynamic regulations of the secretome induced by chronic TNFα, we selected all proteins that were significantly different between TNFα treated and control, and showed a Log2(Fold Change) ≥ |0.38| (Figure S2B). We found 67 proteins to be significantly regulated, on each timepoint, with an overlap of 43 proteins, that were consistently found on both treated and control groups (Figure 6A and B).

**Figure 6.**
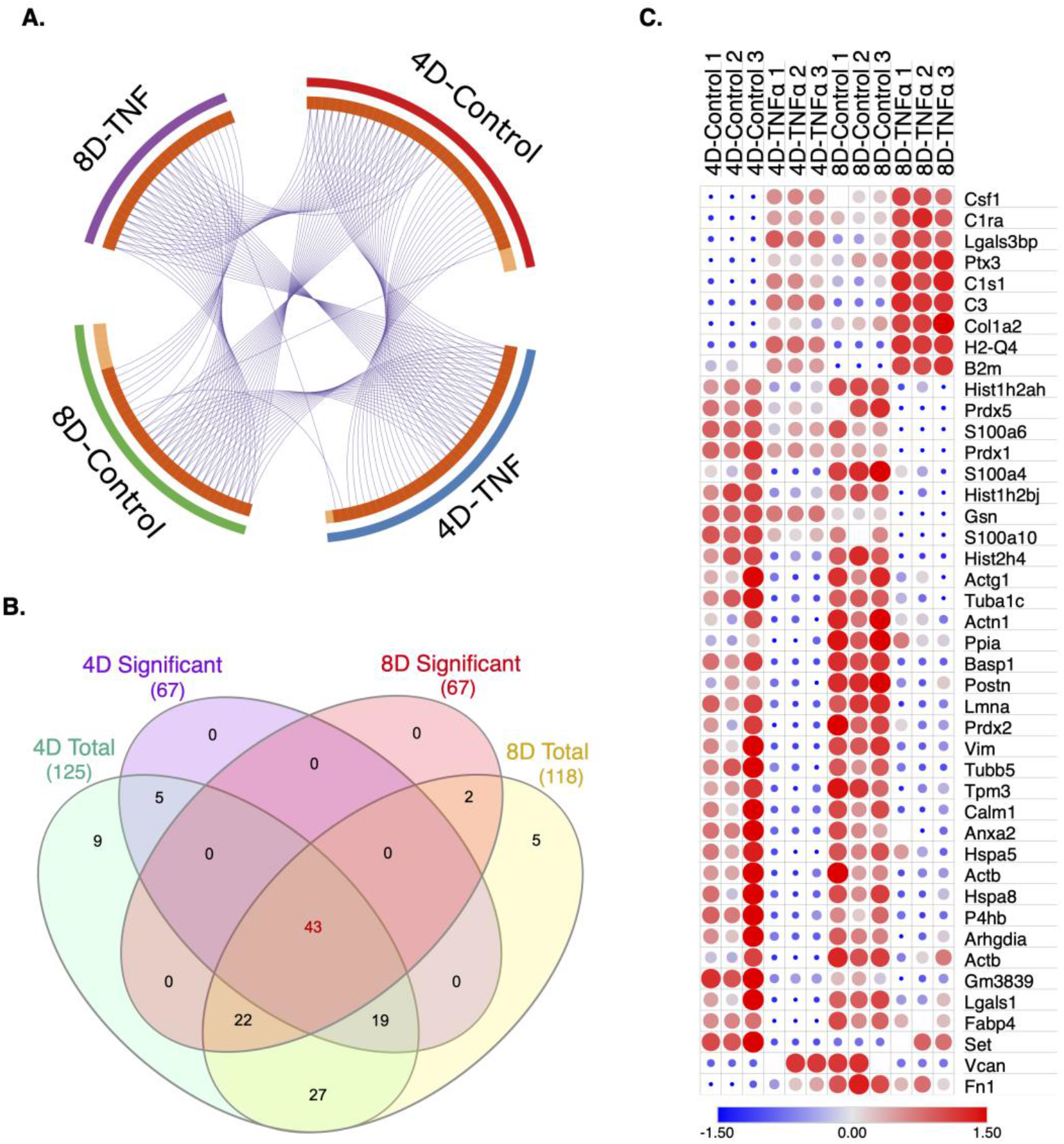
Comparative analysis of 4D and 8D secretomes. (A) Circular plot representation of proteins identified in each treatment group. Light orange indicates uniquely mapped prote ins. Blue strings indicate proteins common to two or more treatment groups. (B) Venn Diagram illustrating the overlap between significantly regulated proteins in 4D (purple) and 8D (green) timepoints. Heat-map depiction of co-secreted proteins between 4D and 8D timepoints Color scale and relative dot size represents Z-scored LFQ values. Blank spaces represent missing values.

In accordance with our global analysis, we focused on the characterization of the proteins that were significantly up or downregulated across 4D and 8D timepoints. Interestingly, our heatmap demonstrates that, from the 43 common proteins, 25 were shown to be downregulated, compared to 7 upregulated, and 3 reversed proteins (Figures 6B). All continuously upregulated proteins directly or indirectly play a role in the progression of inflammatory responses. Worthy of note, all show a clear increase in expression levels at 8D, compared to 4D.

Notably, Galectin-3-binding protein (Lgals3bp) is a protein whose expression is mediated by the canonical activation of the activation of the NF-κB signaling pathway, upon TNFα stimulus (Noma et al., 2012). Lgals3b has been previously demonstrated to induce increased secretion of IL-6 in several different cell lines (Loimaranta et al., 2018), and also promote monocyte migration (Sano et al., 2000). Our results show that TNFα not only promotes the release of Lgals3bp in adipocytes, but its secretion levels increase with prolonged TNFα treatment.

Complement proteins play major roles in acute inflammation, promoting vascular rearrangement, chemotaxis and extravasation of leukocytes (Markiewski and Lambris, 2007). Both Complement C1r-A subcomponent (C1ra) and Complement C3 (C3), constituents of the complement system, were highly upregulated in TNFα treated adipocytes. We also identified a protein, Pentraxin-related protein PTX3 (Ptx3), which modulates the activation of the component system (Daigo et at., 2016), shown to be highly upregulated in our data, suggesting the dysregulation of the component system is likely a key player in the endocrine mediated development of insulin resistance. Collagen alpha-2 (I) chain (Col1a2) was also consistently upregulated in our secretome analysis. Our data supports previous reports demonstrating the overexpression of mRNA levels of collagens types I, III, V, and VI in white adipocytes of diabetic mice (Lin et al., 2016).

In the downregulated protein list, we have identified proteins that alleviate chronic inflammation or are directly suppressed by TNFα. Notably, the secretion of Vimentin (Vim) was particularly downregulated in TNFα treated cells. Vimentin has been shown to attenuate oxidative stress and inflammation in macrophages (Håversen at el., 2018). The downregulation of Vim observed in our secretome analysis, therefore, could implicate enhancement of macrophage induced inflammation.

Along similar lines, galectin-1 (Lgals1) attenuates acute inflammation in rats, and have also been shown to inhibit 3T3-L1 adipocyte differentiation (Rabinovich et al., 2000; Mendez-Huergo et al., 2019). The downregulation of Lgals1 is possibly an important factor that contributes to the dysregulation of adipocyte and macrophage inflammatory responses upon chronic TNFα exposure.

### Phosphoproteome Analysis of Adipocytes at 8D

The insulin signaling cascade is primarily regulated by phosphorylations, and disruptions in phosphorylation patterns are directly associated with the development of insulin resistance (Boucher et al., 2014). To dissect the changes on the cellular phosphoproteome landscape during prolonged (8D) TNFα treatment, we combined our global proteomics with a phosphoproteomic analysis. In short, digested peptides were enriched using the highly selective Polymer-based Metal-ion Affinity Capture (PolyMAC) spin tip (Tymora Analytical), followed by LC-MS/MS analysis (Figure 1SB). Data were searched and analyzed with MaxQuant and Perseus, respectively.

We identified a total of 3134 phosphopepteides 4997 phosphosites, and 1526 phospoproteins. It is important to note that the majority of identified proteins had an LFQ intensity value of zero, in all three replicates, in tandem with a low number of MS/MS counts. The LFQ intensity value is calculated in an extracted ion chromatogram (XIC) by Gaussian fitting 3 or more neighboring datapoints, corresponding to a specific m/z. Thus low abundant peptides are often not quantified as LFQ intensity in the XIC approach. To narrow our analysis to the higher confidence phosphopetides, and to perform a more accurate quantitative comparison between samples, we focus on those proteins and phosphopeptides that were quantified in at least two biological replicates, as previously performed, identifying a total of 113 proteins. By directly comparing the total quantified proteins identified across 8D global proteome, secretome and phosphoproteome analysis, we found 20 proteins present in all three analysis (Table S1), 22 proteins unique to phosphoproteomics, 53 unique to secretomics and 2077 to global proteomics (Figure 7A). The discrepancy between the IDs of identified proteins can be primarily attributed to the identification of proteins with lower abundance in both phosphoproteome and secretome analysis, due to phosphopeptide enrichment and reduced sample complexity, respectively.

**Figure 7.**
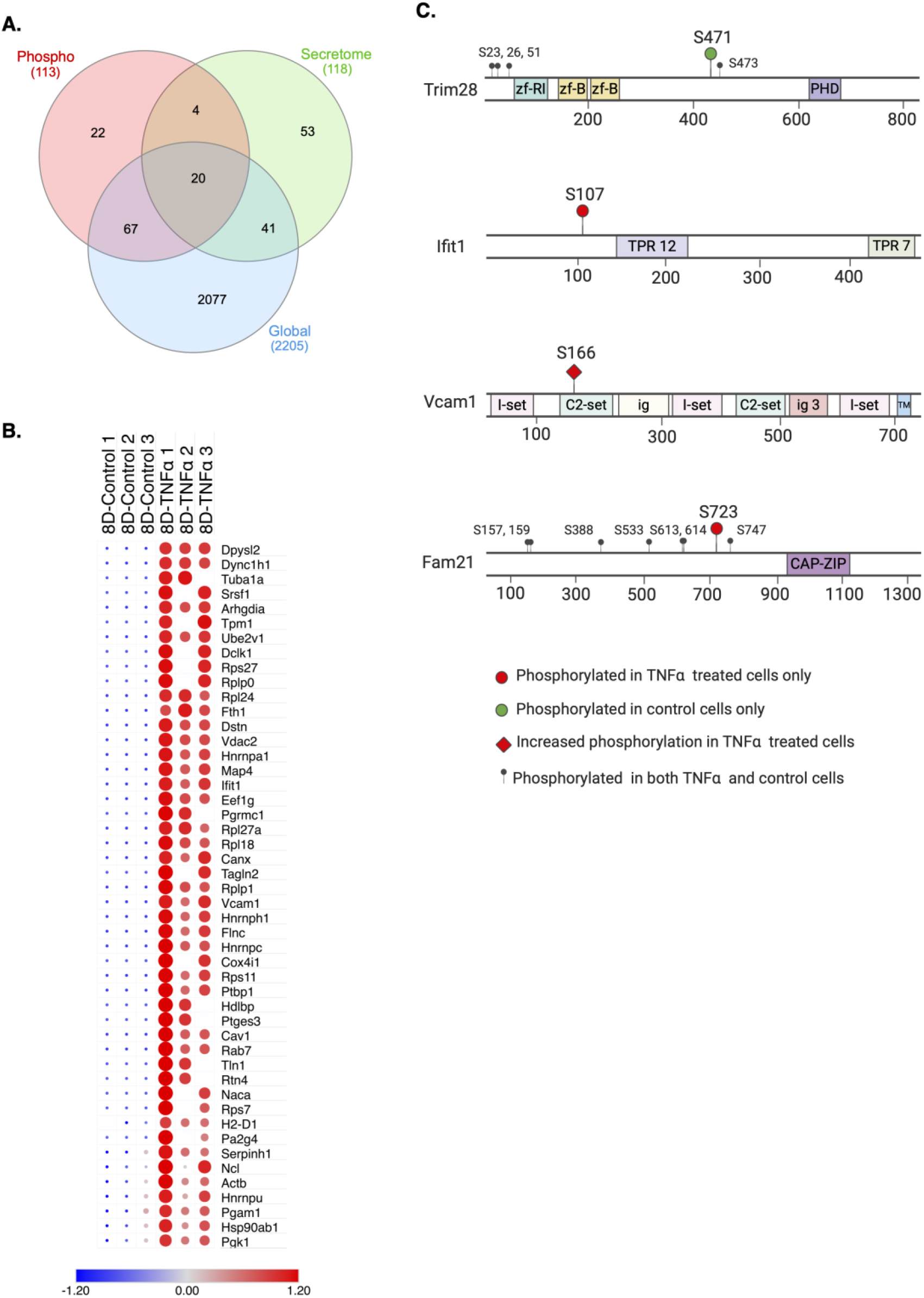
Comparative analysis of 8D phosphoproteomics. (A) Venn Diagram illustrating the overlap between significantly regulated proteins in phosphoproteomic (red), secretomic (green) and global (blue) analysis at 8D. (B) Heat-map depiction of significantly phosphorylated proteins at 8D timepoint. Color scale and relative dot size represents Z-scored LFQ values. Blank spaces represent missing values. (C) Phosphorylated residues identified by MS/MS. Phospho-sites mapped to TNFα treated cells only (red circle), hyperphosphorylated in TNFα treated cells (red diamonds) and control cells only (green circle).

To gain an insight on the changes in phosphorylation levels and patterns upon prolonged TNFα exposure, we utilized the LFQ values of each protein as a direct assessment of phosphorylation. Statistical analysis revealed 48 significantly phosphorylated proteins, all of which were hyperphosphorylated in the TNFα treated group (Figure 7B). Contrary to our expectations, the significant phosphorylated proteins were not directly involved in the insulin signaling pathway, but, remarkably, were primarily downstream effectors of pro-inflammatory cytokines and cytoskeleton-related proteins. One of such proteins, Vascular adhesion protein 1 (Vcam1), which typically shows elevated expression levels in epithelial cells upon TNFα stimulus via NF-κB pathway (Adams and Shaw, 1994; Mullan et al., 2006), was highly phosphorylated in all three TNFα treated biological replicates. Vcam1 has also been previously reported to be upregulated in the membrane fraction of adipocytes treated with TNFα, agreeing with the results here presented (Parker et al., 2016). Another noteworthy protein identified was Interferon-induced protein with tetratricopeptide repeats 1 (Ifit1), an IFNα/β-stimulated genes that has implications primarily in viral and bacterial responses, but have also been shown to stimulate the expression of genes in diverse processes, including apoptosis (Der et al., 1998; Fensterl and Sen, 2015),

The site of phosphorylation in a protein induces a change in the biochemical properties of the residue, and often leads to specific changes in protein function (Ardito et al., 2017). To study the changes in the phosphorylation patterns regulated by prolonged exposure to TNFα, we selected four proteins of interest and identified differentially phosphorylated residues, based on the raw intensity values of the phosphopeptide in each biological replicate (Figure7C). Tripartite motif (Trim) proteins are a family of proteins that primarily function as E3 ubiquitin ligases, and are direct constituents of innate immune response pathways (Rajsbaum et al., 2014). Trim28, particularly, has been reported to act as SUMO E3 ligase, promoting the specific SUMOylation of IFN regulatory factor 7 (IRF7), a key regulator protein of type I IFN-dependent immune response (Liang et al., 2011). We have identified 5 different phosphosites in Trim28, from which one peptide, phosphorylated at Ser 471, was only found in two out of three control replicates (intensity > 0) and was not detected in all TNFα treatment. This might possibly suggest that Ser 471 is uniquely dephosphorylated during prolonged TNFα treatment, and likely implicate a different regulation of IFN-dependent immune response pathways upon TNFα exposure. Taken together with the additional IFN stimulated proteins identified in the global proteomics, these findings further suggest a crosstalk between TNFα and IFNα/β pathways upon chronic TNFα treatment.

Our results also indicate that Ifit1 was uniquely phosphorylated at Ser 107, and Vcam1 was hyperphosphorylated at Ser 166, a phosphorylation site that is here reported for the first time. We have also found that WASH complex subunit FAM21 (Fam21), a protein that has recently been suggested to directly interact with several NF-κB pathway components, and acts as a regulator for p65 binding to IL-6 and IL-1α after TNFα exposure (Deng et al., 2015). Fam21 was uniquely phosphorylated at Ser 723 in chronically treated TNFα cells, indicating the potential involvement of this residue in the activation of Fam21, and subsequent upregulation of NF-κB pathway activation.

Cytoskeleton proteins, such as Tubulin alpha-1A chain (Tuba1a), and cytoskeleton regulator proteins, particularly Filamin-C (Flnc) and Transgelin-2 (Tagln2), both actin binding proteins that have been demonstrated to bind actin and respectively stimulate actin depolymerization and gelation, promoting cytoskeleton rearrangement (Nishida et al., 1985; Shapland et al., 1993), evidencing TNFα involvement in promoting adipocyte restructuring and subsequent macrophage infiltration.

## Discussion

Obesity is a prominent disease, that is often accompanied by chronic adipose tissue inflammation, succeeded by eventual insulin resistance. The development of insulin resistance is tightly linked with the dysregulation of the endocrine and paracrine functions of the adipose tissue, characterized by the hypersecretion of pro-inflammatory adipokines, accompanied by extensive changes in adipocyte cellular landscape (Kahn et al., 2006). The dysfunctional secretion of adipokines, combined with an increased number of necrotic cells culminates in leukocyte invasion, and consequent aggravation of the inflammatory response. The molecular mechanisms underlying insulin resistance are not only complex, but also tightly interconnected. In this study, we unveiled the changes of the adipocyte proteome and phosphoproteome during chronic TNFα treatment, by combining time resolved global proteomic, secretomic and phosphoproteomic analysis.

Firstly, we demonstrated the extensive regulation of protein expression induced by chronic TNFα exposure. Our results indicate a considerably larger dysregulation in the proteome of adipocytes upon chronic TNFα treatment, compared to treatments at shorter timepoints (Chan et al., 2019). These drastic differences in the proteome of TNFα treated adipocytes are likely due to the development of insulin resistance, which is achieved after 4 days of continuous TNFα exposure. The high degree of similarity between the cellular proteome at 4D and 8D further suggests that TNFα promotes sustained changes in protein regulation after the insulin resistance phenotype is developed. Particularly, we have shown that over half of the proteins significantly regulated by TNFα treatment were not only identified in both 4D and 8D timepoints, but also showed very consistent fold change values, indicating the TNFα induces constitutive regulation of proteins that are likely directly involved in pathological progression of insulin resistance. Indeed, we demonstrated that the specific proteins which showed consistent fold change levels across timepoints were directly involved in biological process that drive metabolic disease progression and metabolic inflammation. The downregulation of ceramide catabolism, and upregulation of cytokine secretion and regulation of leukocyte mediated immunity were of special interest. By analyzing the significantly regulated proteins identified and stablishing the processes they partake in, we were able to gain important insight of the crosstalk between different processes, while gaining a holistic perspective of the persistent effects of chronic TNFα exposure.

Our results also demonstrated that chronic TNFα induces prolonged regulations on both the secretome and phosphoproteomics. When combined, our data shows strong correlation between the intracellular, secreted and phosphorylated proteins that are regulated in response to TNFα. Of special importance, we show the significant upregulation of the NF-κB p100/p52 subunit (Nfkb2) in adipocytes chronically treated with TNFα. Nfkb2 processing form p100 to p52 is activated by the non-canonical NF-κB signaling, leading to p52 heterodimerization with RELB and subsequent transcription activation (Zarnegar et al., 2008). The non-canonical NF-κB signaling pathway has been shown to be primarily mediated by TNF receptors (TNFR), but studies have suggested that M-Csf1 receptor (MCSF1R) can also promote the induction of the noncanonical NF-κB signaling during macrophage differentiation (Jin et al., 2014). Interestingly, we have shown that M-Csf1 is highly upregulated in TNFα treated cells, suggesting that the upregulation of M-Csf1 could contribute in the activation of pro-inflammatory responses, activated through the non-canonical NF-κB signaling pathway.

The activation of the canonical NF-κB pathway can also be mediated by TNFR (Beg and Baltimore., 1996), TLR (Kawai and Akira, 2007) and IL-1R (Sanz et al., 2000). Upon activation, downstream heterodimerization of subunits p65 and P50 stimulate the transcription of pro-inflammatory and pro-survival genes. One gene product regulated by the NF-κB pathway is galectin-3-binding protein. Lgals3bp is a secretory glycoprotein, which directly interacts with galectin 1, 3 and 7 (Lgals1, 3, and 7) (Inohara et al., 1996; Lin et al.; 2015). Although Lgals3 and Lgals3bp have been principally linked to tumor aggressiveness and metastasis (Takenaka et al., 2002; Natarajamurthya et al., 2019), previous findings have suggested Lgals3 role in macrophage recruitment (Sano et al., 2000). The high upregulation of Lgals3bp we observed in both global proteomics and secretomics of TNFα treated adipocytes shed light on the potential pro-inflammatory action of Lgals3 and Lgals3bp. Our results are the first to demonstrate the upregulation of Lgals3bp induced by TNFα in adipocytes, and is a potential candidate for further mechanism elucidation. Interestingly, our results have also shown another galectin, Lgals1, to be significantly downregulated in the secretome of adipocytes treated with TNFα. Lgals1 has several anti-inflammatory properties, including the inhibition proinflammatory cytokine synthesis, leukocyte migration, degranulation and survival (Sundblad et al., 2017), which further suggest potential important roles played by the Galectins and Galectin-3-binding protein on the progression of metabolic inflammation.

Toll like receptors 2, 3 and 4 (TLR) mediate the NF-κB pathway activation, which leads to the transcription activation of pro-inflammatory cytokines, including IFNβ (Kawai and Akira, 2007). We have here shown a high and consistent upregulation of global TLR2, accompanied by the regulation of several interferon-induced protein with tetratricopeptide repeats, such as Ifit2 and Ifit3, identified in the global proteomics and hyperphosphorylated Ifit1, identified on the phosphoproteomics. Although the exacerbation of metabolic inflammation and TNFα expression in adipose tissue by IFN cytokines, secreted by T-cells, has been previously demonstrated (Rocha et al., 2008), our results suggest a potential crosstalk of the NF-κB signaling pathway and IFN-family cytokine secretion, induced by chronic TNFα exposure, occurs in adipocytes. This crosstalk is also supported by the activation of TLR2 by ceramides (Majumdar et al., 2002). Our data evidences a downregulation of the ceramide catabolic biological process, which likely implicates the excess accumulation of ceramides. Accordingly, the overexpression of Asah1 has been shown to attenuate the inhibitory effects of free fatty acids in insulin sensitivity in myotubes (Chavez et al., 2005). The downregulation of Asah1 mediated by TNFα, combined with the downregulation of Psap and Naga, likely induces increase in the concentration of free fatty acids, which, in turn, activate TLR2, enhancing inflammation.

Taken together, our data suggest an overall upregulation of the canonical and non-canonical activation of the NF-κB pathway, mediated by TNFR and TRL, inducing the regulation of several proteins, that culminate in pro-inflammatory responses and insulin signaling disruption. We have identified key proteins expressed in adipocytes that have previously only been associated with immune cells, suggesting that adipocytes could ectopically express and secrete additional cytokines, that can be potential targets for future therapeutic strategies.

## Materials and methods

### Cell culture

Murine 3T3-L1 (CL-173™) pre-adipocytes (ATCC, Manassas, VA, USA) were cultured in DMEM (ATCC) supplemented with 10% bovine calf serum (ATCC). Chemically-induced differentiation was performed following the supplier’s standard protocol (ATCC). Briefly, differentiation was induced for 48 hours by differentiation media treatment (DMEM supplemented 10% Fetal Bovine Serum (FBS), 1.0 μM Dexamethasone, 0.5 mM Methylisobutylxanthine (IBMX), 1.0 μg/mL Insulin and 1 uM Rosaglitazone). Differentiation media was subsequently replaced with Adipocyte Maintenance Medium (DMEM supplemented with 10% Fetal Bovine Serum 1.0 μg/mL Insulin), and cells were further cultured in Adipocyte Maintenance Medium for 6 days. Adipocytes were chronically treated with 10 nM insulin, and experimental group were treated with an additional 2ng/mL TNFα, for a total of 4 or 8 days. Treated media was replaced with Serum Free media with or without TNFα for 12 hours. Cells were then treated with 10 nM insulin for 30 minutes, followed by spent media and cell collection.

### Cell lysis and protein extraction

Collected cells were washed three times with cold 1X PBS and subsequentially resuspended in 100 mM ABC supplemented with protease and phosphatase inhibitors. Cell suspension was transferred to precellys homogenization vials (Bertin Technologies SAS, France), and samples were homogenized for 90 seconds at 6500 rpm. Protein concentration was measured by bicinchoninic acid (BCA) assay (Pierce Chemical Co., Rockford, IL, USA). 300 ug of total protein were precipitated with 4 volumes of cold acetone at −20 °C overnight. Samples were centrifuged at 13500 RPM for 10 minutes, acetone supernatant was removed, and pellets were dried in a vacuum centrifuge for 15 minutes.

### Buffer exchange and media concentration for secretome analysis

2 mL of collected spent media was loaded in Vivaspin 6™ Centrifugal Concentrator 10,000 MWCO PES (Sartorius AG, Germany) and centrifuged at 4000 RPM for 13 minutes to concentrate samples. 2 mL of 25 mM ABC were then added to each tube, followed by centrifugation at 4000 RPM for 13 minutes, for a total of three washes. Final protein concentration was measured by BCA assay, and 50 ug were precipitated with 4 volumes of cold acetone at −20 °C overnight. Samples were centrifuged at 13500 RPM for 10 minutes, acetone supernatant was removed, and pellets were dried in a vacuum centrifuge for 15 minutes.

### Sample preparation for MS analysis

Dried samples were fully resuspended in 60 uL 8 M urea, 10 mM DTT and incubated at 37 °C for 1 hour. 60 uL of alkylation reagent mixture (97.5% Acetonitile (ACN), 0.5% Triethylphosphine, 2% Iodoethanol) was added to samples, which were again incubated at 37 °C for 1 hour. Samples were dried in a vacuum centrifuge and resuspended in 200 uL of 0.05 ug/uL Lys-C/Trypsin (Promega, WI, USA) in 25 mM Ammonium Bicarbonate. Proteolysis was carried out using a barocycler (50°C, 60 cycles: 50 s at 20 kPSI and 10 s at 1 ATM). Digested samples were desalted with Pierce Peptide Desalting Spin Columns (Thermo Fisher Scientific, IL, USA). Phosphopeptide enrichment was subsequently performed using PolyMac spin tips (Tymora Analytical, IN, USA), following manufacturer’s recommendations.

### Mass spectrometry

Samples were resuspended in 3% ACN, 0.1% Formic Acid (FA) and separated by reverse-phase HPLC with an Acclaim PepMap 100 C18 analytical column (75 μm ID × 50 cm) packed with 2 μm 100 Å PepMap C18 medium (Thermo Fisher Scientific), coupled with the Q-Exactive Orbitrap HF (Thermo Fisher Scientific) mass spectrometer. Peptides for global analysis were separated in a 160 minute gradient. Peptides were loaded into the column with 2% mobile phase solution B (80%ACN with 0.1% FA in water). Solution B was linearly increased to 27% B until 110 minutes, followed by an increase to 40% B at 125 minutes. Mobile phase solution B was subsequentially increased 100% at 135 minutes, and held constant for an additional 10 minutes. The gradient was then returned to 2% B until the end of the run. Phosphopetideas and peptides for secretome analysis were separated in a 120 minute gradient. Peptides were loaded into the column with 2% mobile phase B. Mobile phase solution B was increased linearly to 30% B until 80 minutes, followed by an increase to 45% B at 91 minutes. Mobile phase solution B was subsequentially increased 100% at 93 minutes, and held constant for an additional 5 minutes. The gradient was then returned to 2% B until the end of the run. Data-dependent acquisition MS/MS was performed for the top 20 precursors, with MS/MS spectra recorded from 400 to 1600 m/z.

### Data analysis

Search of raw MS/MS data was performed using MaxQuant software against Uniprot *Mus musculus* database. The search was performed using Lys-C/Trypsin enzymes for specific digestion, at 2 missed cleavage allowance. Methionine oxidation, and for phosphoproteomics STY phosphorylation, were set as variable modifications and Iodoethanol set as fixed modification. Main search peptide tolerance was set to 10 ppm, and false discovery rate (FDR) of peptides and proteins identification was set to 1%. Peptide quantitation was performed using “unique plus razor peptides”. The resulting MaxQuant data was analyzed with Perseus platform. “Contaminants”, “reverse”, and “only identified by site” proteins were filtered out, and LFQ intensity values were Log2 transformed. Proteins were then filtered based on 2 minimum valid values in at least one group, and MS/MS count ≥ 2 in a least two replicates. Missing values were zero-filled with the value corresponding to half the minimum LFQ intensity value. Statistical analysis was performed using a two tailed Student’s t test. Proteins with a P value ≤ 0.05 and absolute Log_2_(LFQ) ≥ |0.38| were considered significantly regulated. Gene ontology (GO) was done using Metascape online software, with only Biological Processes (GO) selected for annotation, membership and enrichment.

### Western blotting

20 ug of cell lysate were combined with NuPAGE™ LDS Sample Buffer (4X) (Thermo Fisher Scientific) and water, heated at 70°C for 10 minutes, and subsequent western blot was performed as previously described (Zheng et al., 2014). The following antibodies and dilutions were used: M-CSF Polyclonal Antibody (Thermo Fisher Scientific, catalog # PA5-42558, RRID AB_2609577, 1:500), MIF Polyclonal Antibody (Thermo Fisher Scientific, catalog # PA5-82117, RRID AB_2789278, 1:500), Monoclonal Anti-β-Actin Antibody (Sigma-Aldrich, catalog #A5441, 1:10000). Secondary antibodies and dilutions used were: Goat anti-Rabbit IgG (H+L) Highly Cross-Adsorbed Secondary Antibody, Alexa Fluor 680 (Thermo Fisher Scientific, catalog # A-21109, RRID AB_2535758, 1:10000) and Goat anti-Mouse IgG (H+L) Cross-Adsorbed Secondary Antibody, Alexa Fluor 790 (Thermo Fisher Scientific, catalog # A11375, RRID AB_2534146, 1:1000). Blots were visualized with the Odyssey® CLx Imaging System.

## Supporting information

SI Table 1: 4D and 8D Total Protein List from Global Proteomics

SI Table 2: 4D and 8D Total Peptide List from Global Proteomics

SI Table 3: 4D Significantly Regulated Protein List from Global Proteomics

SI Table 4: 8D Significantly Regulated Protein List from Global Proteomics

SI Table 5: 4D Total Protein List from Secretomics

SI Table 6: 8D Total Protein List from Secretomics

SI Table 7: 4D Total Peptide List from Secretomics

SI Table 8: 8D Total Peptide List from Secretomics

SI Table 9: 4D Significantly Regulated Protein List from Secretomics

SI Table 10: 8D Significantly Regulated Protein List from Secretomics

SI Table 11: 8D Total Protein List from Phosphoproteomics

SI Table 12: 8D Total Peptide List from Global Phosphoproteomics

SI Table 13: 8D Significantly Regulated Protein List from Global Phosphoproteomics

## Acknowledgments

This work was partially supported by grants from the Indiana Clinical and Translational Science Institute (CTSI). All the LC-MS/MS experiments were performed at the Purdue Proteomics Facility in the Bindley Bioscience Center. We thank Dr. Tiago Sobreira for invaluable discussions regarding the statistical analysis and data interpretation. We also kindly thank Dr. Wen H. Wang for proving access to his western blot equipment and Dr. Andrisani Lab at the College of Veterinary Medicine at the Purdue University for actin antibody. Illustrations were created with BioRender.com.

## Author contributions

Conceptualization, R.M. and U.K.A.; Methodology, R.M. and U.K.A; Cell culture, treatment and sample preparation, R.M.; LC-MS/MS data acquisition, U.K.A., Data analysis, R.M., Writing – Original Draft, R.M.; Writing – Review & Editing, U.K.A.; Fund Acquisition, U.K.A, Supervision, U.K.A.

## Data availability

All Raw LC-MS/MS data are deposited in the MassIVE data repository (massive.ucsd.edu/), with ID: MSV000085562

---

**Figure S1.**
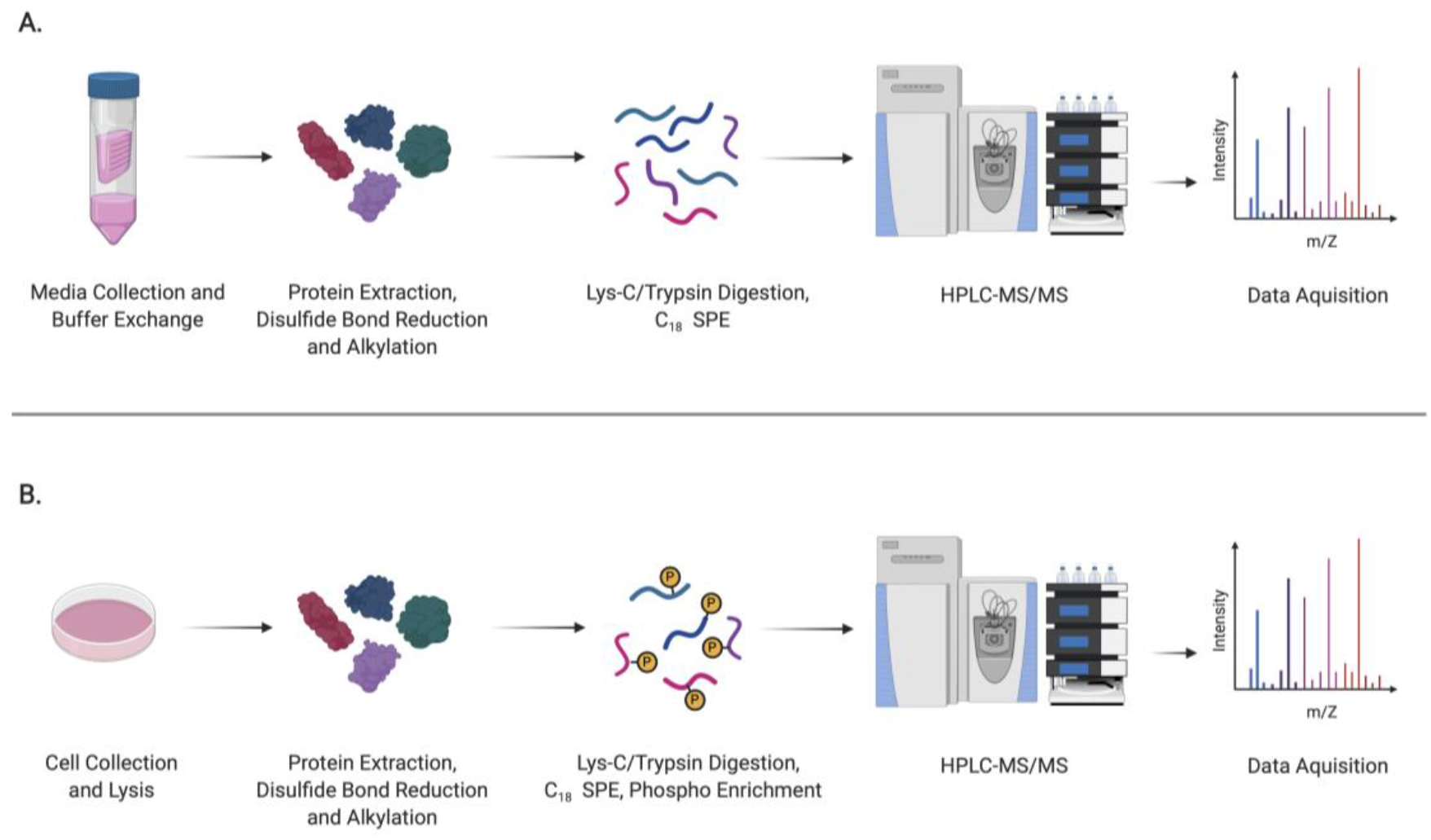
Experimental design and label-free quantitative secretomic and phosophoproteomic analysis workflow. (A) Spend media was collected and concentrated. Protein was extracted, with sequential reduction and alkylation of cysteine disulfide bonds. Proteins were digested with Lys-C/Trypsin, desalted and analyzed by HPLC-MS/MS. (B) Protein was extracted, with sequential reduction and alkylation of cysteine disulfide bonds. Proteins were digested with Lys-C/Trypsin, desalted, and phosphopeptides were enriched before HPLC-MS/MS analysis.

**Figure S2.**
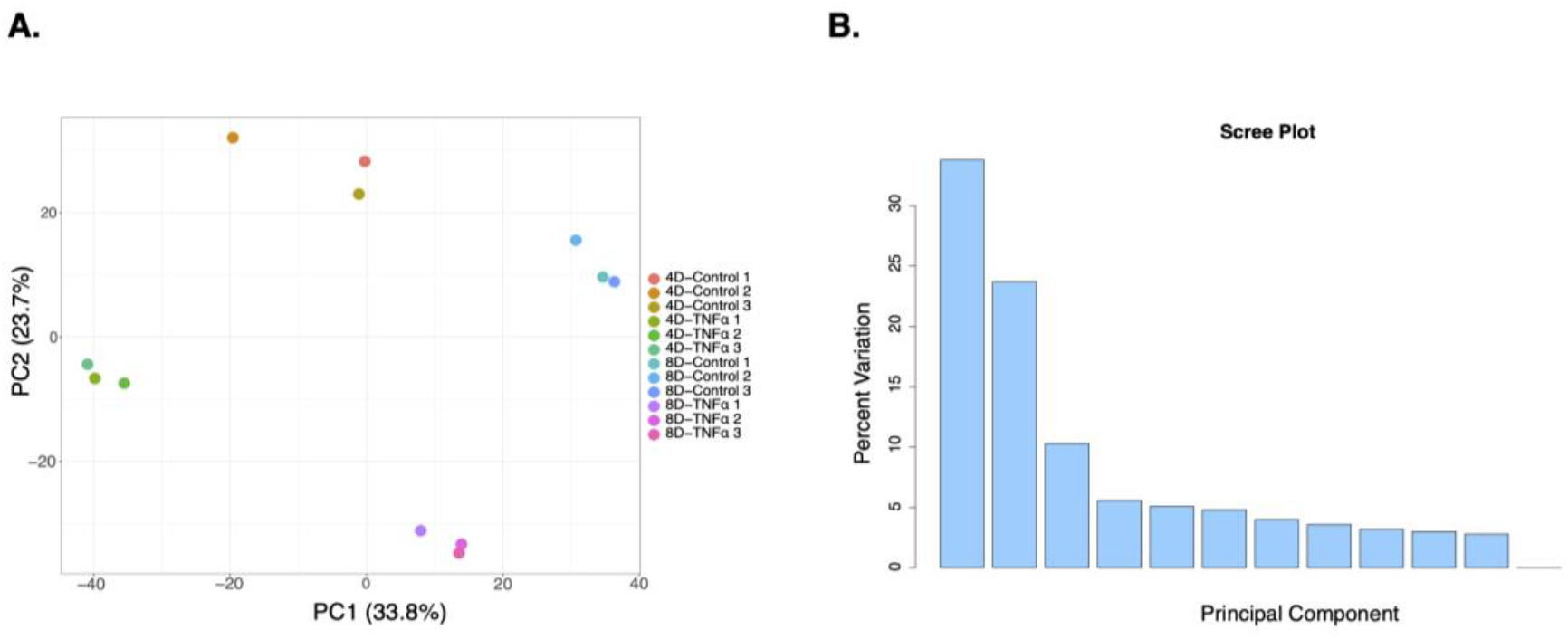
Treatment groups are highly correlated: (A) PCA plot depiction of all replicates analyzed in the global proteomics. Individual samples are indicated by color, indicated in the figure legend. (B) Scree plot representation of Principal Components.

**Table S1:** Commonly identified proteins in 8D global proteome, secretome, phosphoproteome.

| Protein IDs | Protein names | Gene names |
| --- | --- | --- |
| Q5FWJ3 | Vimentin | Vim |
| P16045 | Galectin-1 | Lgals1 |
| B2RTM0 | Histone H4 | Hist2h4 |
| Q8VDD5 | Myosin-9 | Myh9 |
| P48678 | Prelamin-A/C | Lmna |
| P09411 | Phosphoglycerate kinase 1 | Pgk1 |
| Q71LX8 | Heat shock protein HSP 90-beta | Hsp90ab1 |
| Q3UGY5 |  | Fn1 |
| Q544Y7 | Cofilin-1 | Cfl1 |
| P99024 | Tubulin beta-5 chain | Tubb5 |
| Q64727 | Vinculin | Vcl |
| Q3U7Z6 | Phosphoglycerate mutase 1 | Pgam1 |
| Q5EBQ2 | Phosphatidylethanolamine-binding protein 1 | Pebp1 |
| Q99PT1 | Rho GDP-dissociation inhibitor 1 | Arhgdia |
| Q9QZ83 |  | Actg1 |
| P63101 | 14-3-3 protein zeta/delta | Ywhaz |
| Q3TZQ2 | Moesin | Msn |
| Q9WVA4 | Transgelin-2 | Tagln2 |
| P08228 | Superoxide dismutase [Cu-Zn] | Sod1 |
| Q91XV3 | Brain acid soluble protein 1 | Basp1 |

## Supplementary information

**SI Table 1: 4D and 8D Total Protein List from Global Proteomics**

**SI Table 2: 4D and 8D Total Peptide List from Global Proteomics**

**SI Table 3: 4D Significantly Regulated Protein List from Global Proteomics**

**SI Table 4: 8D Significantly Regulated Protein List from Global Proteomics**

**SI Table 5: 4D Total Protein List from Secretomics**

**SI Table 6: 8D Total Protein List from Secretomics**

**SI Table 7: 4D Total Peptide List from Secretomics**

**SI Table 8: 8D Total Peptide List from Secretomics**

**SI Table 9: 4D Significantly Regulated Protein List from Secretomics**

**SI Table 10: 8D Significantly Regulated Protein List from Secretomics**

**SI Table 11: 8D Total Protein List from Phosphoproteomics**

**SI Table 12: 8D Total Peptide List from Global Phosphoproteomics**

**SI Table 13: 8D Significantly Regulated Protein List from Global Phosphoproteomics**

